# Star-Shaped Glatiramer Acetate Mitigates Pulmonary Dysfunction and Brain Neurodegeneration in a Murine Model of Cryptococcus-Associated IRIS

**DOI:** 10.1101/2025.01.07.631707

**Authors:** Shehata Anwar, Jinyan Zhou, Lauren Kowalski, Joshua Saylor, Devanshi Shukla, Katelyn Boetel, Ziyuan Song, Kamal Sharma, Jianjun Cheng, Makoto Inoue

## Abstract

Cryptococcal-associated immune reconstitution inflammatory syndrome (C-IRIS) is a clinical worsening or new presentation of cryptococcal disease following the initiation of antiretroviral therapy. C-IRIS is primarily driven by an influx of pathological CD4^+^ T cells, which triggers a hyperinflammatory response. The murine model of C-IRIS is a way to study the disease in mice and understand how the immune system triggers life-threatening outcomes in patients. We previously developed a murine C-IRIS model and demonstrated that C-IRIS is triggered by pathological CD4^+^ T cells, particularly Th1 cells, in the brain, which triggers neurodegeneration and pulmonary dysfunction. Using this unique mouse model, we tested the therapeutic effect of a star-shaped glatiramer acetate (sGA), which is a more effective isomeric form than linear GA. Here, we observed that sGA suppresses Th1 differentiation in the lung tissues, reducing CD4^+^ T cell and Th1 cell count. It also reduced microglia populations in the brain. Together, these changes improved respiratory dysfunction caused by C-IRIS, lowered mortality rate, and reduced brain neurodegeneration. These findings suggest that sGA could be an effective therapeutic strategy for managing C-IRIS.

## Introduction

*Cryptococcu*s-associated immune reconstitution inflammatory syndrome (C-IRIS) occurs when a patient who has been infected with *Cryptococcus neoformans* (Cn) has an overactive immune system during the process of reconstitution having proinflammatory responses as the immune system recovers and is in danger of death1C-IRIS is a common complication, with approximately 25-30% of HIV patients with cryptococcal infections developing IRIS within the first four months of combination antiretroviral therapy (cART), with mortality rates varying widely but may exceed 50%(Sereti et al., 2020). Patients with C-IRIS can exhibit neurological symptoms such as headaches, fever, changes in mental status, and memory loss. Additionally, they may show non-neurological symptoms like pulmonary disease(Lortholary et al., 2005; Skiest et al., 2005; Haddow et al., 2010; Perfect et al., 2010; Thambuchetty et al., 2017; Hu et al., 2020). Nonsteroidal anti-inflammatory drugs (NSAIDs) and corticosteroids are prescribed to suppress excessive inflammation in C-IRIS patients(Meintjes et al., 2012a). Such immunosuppressive treatments may impair response to the existing infection and increase susceptibility to new infections(Hsu et al., 2011; Tani et al., 2012). Therefore, understanding the immunopathogenesis of C-IRIS to facilitate targeted interventions is critical.

Patients with C-IRIS typically present with a high level of circulating CD4^+^ T cells, particularly of the Th1 subtype(Boulware et al., 2010; Neal et al., 2017). Recently, we have developed a mouse model of C-IRIS using immunocompromised *Tcra^-/-^*mice, which lack T cells, or *Rag1*^-/-^ mice, which lack T and B cells, with intranasal (i.n.) infection of Cn serotype A H99 (CnH99) and intravenous (i.v.) transfer of CD4^+^ T cells three weeks after CnH99 infection (hereinafter C-IRIS condition/mice)(Khaw et al., 2020; Kawano et al., 2023b). This mouse model showed manifestations of weight loss, high mortality, systemic upregulation of proinflammatory cytokines, elevated levels of CD4^+^ T cells/Th1 in the lungs, infiltration of CD4^+^ T/Th1 cells into the brain, and brain edema(Khaw et al., 2020; Kawano et al., 2023b). We further demonstrated that pulmonary dysfunction associated with the C-IRIS condition in mice could be attributed to brain neurodegeneration by infiltration of CD4^+^ T cells into the brain(Kawano et al., 2023b). Therefore, targeting CD4^+^ T /Th1 cells could be a promising therapeutic target for C-IRIS.

Glatiramer acetate (GA), also known as Copolymer 1 (Cop 1) or Copaxone, is a synthetic amino acid copolymer and disease-modifying therapy (DMT) used to treat multiple sclerosis (MS) (Tselis et al., 2007). It’s an injectable immunomodulator medication approved by the FDA to reduce the frequency of relapses in relapsing and relapsing-remitting MS(Johnson et al., 1998; Tselis et al., 2007). GA has also effectively treated clinically isolated syndrome (CIS) (Prod’homme and Zamvil, 2019). Moreover, early subcutaneous injection of GA in APP/PS1 mice, a mouse model of Alzheimer’s disease, delayed disease progression, reduced amyloid-β plaque burden, regulated neuroinflammation, promoted neuroprotection, and induced dendritic-like microglia(Huang et al., 2024). GA is thought to work by activating suppressor Th1 cells, interfering with the antigen-presenting function of specific immune cells, and consecutively increasing the secretion of anti-inflammatory cytokines(Schrempf and Ziemssen, 2007). Additionally, GA-specific T helper cells migrate to the brain and suppress neuroinflammation(Aharoni et al., 2003). Modifying the linear random co-polypeptide form into a star-shaped GA (sGA) exhibited much better performance than linear GA in treating experimental autoimmune encephalomyelitis (EAE), a mouse model of MS(Song et al., 2020). Moreover, sGA has prolonged drug retention and faster uptake by antigen-presenting cells, attributed to its spherical structure and larger size(Song et al., 2020). Therefore, we investigated the therapeutic effect of sGA on the experimental mouse model of C-IRIS.

## Materials and Methods

### Mice

C57BL6/J and *Rag1*^-/-^ (002216) mice purchased from The Jackson Laboratory were used for the current work. Mice were group-housed with two to five in an individual cage and kept under specific pathogen-free conditions with a 12-h light/dark cycle. Sterile pelleted mouse diet and water were given ad libitum for mice health monitoring. Healthy male and female mice aged 16-20 weeks were randomly selected for this study. All mouse experiments were approved by the University of Illinois Animal Care and Usage Committee (IACUC) under protocol number 22140.

### C-IRIS induction

*Cryptococcus neoformans* (Cn) serotype A strain H99 (CnH99, ATCC 208821) was used from fresh isolates from frozen stocks without serial plating. CnH99 was cultured on yeast extract-peptone-dextrose (YPD) medium (yeast extract 1%, peptone 2%, dextrose 2%) plates at 30 °C for 48 hours. An isolated CnH99 colony was incubated on a YPD liquid medium at 30 °C with 200 rpm shaking overnight. CD4^+^ T cells were isolated from the spleen and inguinal/axillary lymph nodes of naïve C57BL6 mice (6-8 weeks old). Through the negative selection, cells were labeled with biotinylated antibodies CD19 (Cat# 115504, BioLegend), CD11c (Cat# 117304, BioLegend), CD8 (Cat# 100704, BioLegend), Ly6G (Cat# 127604, BioLegend), and CD11b (Cat# 101204, BioLegend) and streptavidin-coated magnetic particles (STEMCELL, Vancouver, Canada, EasySep™ Streptavidin RapidSpheres™ Isolation Kit, Cat# 19860A). B, CD8^+^ T cells, neutrophils, dendritic cells, and macrophages were separated. Then, through the positive selection using biotinylated CD4 antibody (Cat# 100404, BioLegend) and Biotin-coated magnetic particles (STEMCELL, EasySep™ Biotin Positive Selection Kit, Cat# 17665), CD4^+^ T cells were isolated. *Rag1^-/-^* male mice aged 16-20 weeks old were anesthetized with isoflurane (5% for induction, 3% for maintenance) in oxygen (2 L/min) and intranasally infected with CnH99 (100 yeasts in 30 µl PBS). The C-IRIS model was induced by intravenous (i.v) injection (10^6^ cells in 200 µl PBS with 2% FBS) of isolated CD4^+^ T cells into Cn-infected mice 3 weeks after CnH99.

### Drug injection

sGA was administered intraperitoneally at a dose of 1 mg/kg in a 200 µl vehicle of phosphate-buffered saline (PBS), with two-time dosing on day 1 and day 3 post-CD4^+^ T cell injection (C-IRIS induction). The mice were randomized to vehicle-treated and sGA-treated groups(Song et al., 2020).

### Whole body plethysmography

The experimental mice groups (sGA-injected C-IRIS, vehicle-injected C-IRIS, and naïve mice) were acclimated in the room and placed in the testing chamber under no anesthesia or restraint for recording for 5 min using the Buxco Small Animal Whole Body Plethysmography apparatus (Data Sciences International). The apparatus comprises a calibration unit, a flow generator, and a testing chamber. The testing chamber was cleaned with 70% ethanol and dried between trials. Respiratory parameters were recorded by the device, and the data were analyzed using Ponemah® Software, Ver 5.2.

### Golgi staining

Isoflurane deeply anesthetized mice were perfused with 4% PFA via transcardial perfusion. According to the manufacturer’s instructions, Golgi-Cox staining was done as described by FD Rapid Golgi-Stain Kit (FD Neuro-Technologies, Inc.). Briefly, brains were harvested and transferred into 10 ml of Solution A and B at a ratio of 1:1 for 24 hours, followed by another 14 days in the dark in the same amount of Solution A and B. Brains were then placed in solution C for three days, with a new solution being substituted after 24 hours. Then, brains were embedded in Tissue-Tek OCT compound, coronally cut into 60 µm sections, stored in 0.02% azide solution at 4 C, and mounted on poly-L-lysine coated glass slides. Images were taken using a light microscope and analyzed using Image J software.

### Flow cytometry

Seven days after CD4^+^ T cell transfer, brains and lungs were retrieved. Tissues were excised and minced in PBS supplemented with collagenase D (1 mg/ml). Minced tissues were incubated for 30 minutes at 37°C, filtered through the 80-μm mesh, and centrifuged at 277 *g* at 4°C. To isolate mononuclear cells from the brains and lungs, cells were resuspended in 30% Percoll (in PBS), laid over 70 % Percoll, and centrifuged at 377 *g* for 20 minutes at room temperature. Before cell staining with antibodies, Fc receptors were blocked with Fc Block (BD Pharmingen) for 7 min on ice. Cells were stained with fluorochrome-conjugated specific antibodies (1:200) for 30 mins on ice, followed by washing and collecting cells. Stained cells were analyzed on Cytek Aurora with the FCS Express (De Novo). Antibodies used were: APC-Fire 750 CD45 (Cat# 103154, BioLegend), Pe/Cy5 CD3 (Ref# 15-0031-82, eBioscience), Pe/Cy7 CD4 (Cat# 100422, BioLegend), FITC-IFNγ (Cat# 505806, BioLegend), PE-IL17 (Cat# 506904, BioLegend), and PE CD11b (Cat# 101208, BioLegend). Data were analyzed using FCS Express version 6.

### Statistical Analyses

Statistical analysis was evaluated with two-tailed unpaired Student’s t-tests or Log-rank (Mantel-Cox) test following one-way ANOVA using GraphPad Prism Version 9. Data are presented as mean ± SEM, and *P*-values *P ≤ 0.05 were considered statistically significant. Animals were randomly used for experiments under the criteria aforementioned in the section “Animals.” All behavior experiments were performed in a blinded and randomized fashion. No animals or data points were excluded. Experiments were repeated at least two times. Measurements were taken from distinct samples for each experiment. For each test, the experimental unit was an individual animal. No statistical methods were used to predetermine sample sizes, but our sample sizes are similar to those generally employed in the field(Inoue et al., 2016; Oakley et al., 2016). Data distribution was assumed to be normal, but this was not formally tested. Statistical analyses and graphical presentations were computed with GraphPad Prism software (GraphPad, La Jolla, United States). Where appropriate, the n (number of mice or biological samples) and p values are indicated in the figure legends.

## Results

### sGA suppressed pulmonary dysfunction and reduced mortality in C-IRIS mice

Because pulmonary dysfunction is one of the major clinical signs in C-IRIS patients(Lortholary et al., 2005; Skiest et al., 2005; Haddow et al., 2010; Perfect et al., 2010; Thambuchetty et al., 2017; Hu et al., 2020), we conducted a whole-body plethysmography (WBP) study to evaluate the respiratory dysfunctions, where mice were in an unanesthetized and unrestrained state and their pulmonary functions were recorded. Compared to naïve mice, C-IRIS condition/mice received intranasal (i.n.) infection of Cn serotype A H99 (CnH99) into *Rag 1^-/-^*mice and intravenous (i.v.) transfer of CD4^+^ T cells three weeks after CnH99 infection (abbreviation already explained)(Khaw et al., 2020; Kawano et al., 2023a). Compared to naïve, C-IRIS mice showed pulmonary dysfunctions, such as reduced breaths per minute (BPM) and minute volume (MV), and increased total time (TT) per cycle, expiration time (ET), and inspiration time (IT) at 5 days after CD4^+^ T cells transfer (**Fig. 1**), similar to a previous report(Kawano et al., 2023b). On the other hand, when sGA was administered intraperitoneally (i.p.) at a dose of 1 mg/kg(Song et al., 2020) with two-time dosing on days 1 and 3 after CD4^+^ T cell transfer into CnH99-infected mice (C-IRIS induction), these pulmonary dysfunctions were significantly prevented (**Fig. 1**). Briefly, C-IRIS mice that were injected with sGA showed a significant decrease of alterations of BPM, MV, TT, ET, and IT, compared with vehicle-injected C-IRIS mice.

**Figure 1.**
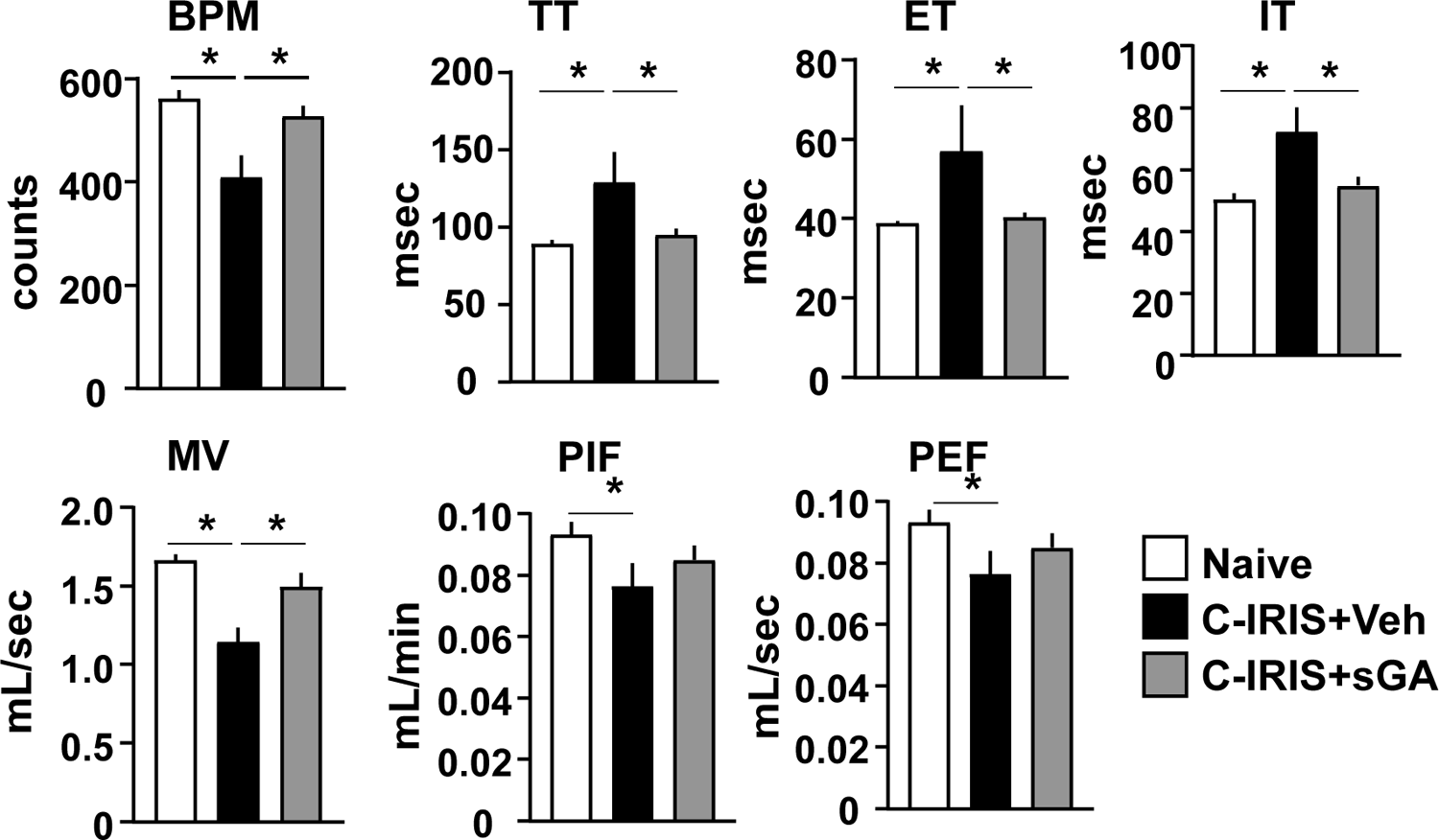
sGA prevents pulmonary dysfunction in C-IRIS mice. Whole-body plethysmography (WBP) assessed pulmonary function in C-IRIS mice. C-IRIS mice exhibited significant respiratory dysfunction, including reduced breaths per minute (BPM), minute volume (MV), and increased total time (TT), expiration time (ET), and inspiration time (IT) compared to naive controls. Treatment with sGA (1 mg/kg, intraperitoneally on days 1 and 3 post-CD4+ T cell transfer) significantly restored normal respiratory function. Data are expressed as mean ± SD (*p < 0.05).

Pulmonary dysfunction in C-IRIS mice correlates with high mortality rates (Khaw et al., 2020). Therefore, survival analysis was conducted in *Rag1^-/-^* mice with C-IRIS, and we found significantly higher mortality rates in C-IRIS mice than the naïve ones (**Fig. 2**). Moreover, sGA treatment (1 mg/kg, days 1 and 3 after CIRIS induction) in C-IRIS mice significantly reduced mortality rates (**Fig. 2**). Together, these results suggest that sGA is an effective treatment for preventing pulmonary dysfunction and mortality due to C-IRIS induction.

**Figure 2.**
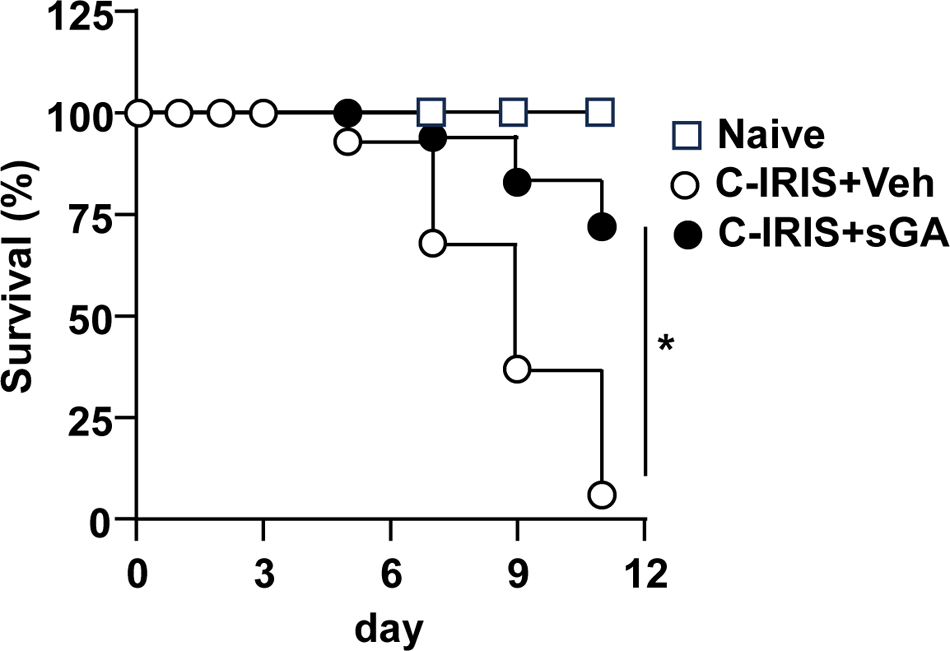
sGA reduces mortality in C-IRIS mice. Survival analysis showed a significantly higher mortality rate in untreated C-IRIS mice compared to naive controls. sGA treatment (1 mg/kg, i.p.) significantly reduced mortality rates. Kaplan-Meier survival curves are presented with significance denoted by (*p < 0.05).

### sGA decreased inflammatory cell populations in the lung and brain of C-IRIS mice

Because it is well established that GA treatment reduces proinflammatory Th cell differentiation, such as Th1 differentiation(Simpson et al., 2002; Kantengwa et al., 2007; Ziemssen and Schrempf, 2007), we tested whether sGA treatment reduces Th1 populations in the lung, a primary infectious site of CnH99. Using flow cytometry analysis, sGA treatment significantly reduced numbers of CD4^+^ T, Th1, and Th17 cells in the lung at 5 days after CD4^+^ T cell transfer in CnH99-infected mice (**Fig. 3A**). Previously, we found that CD4^+^ T cell infiltration into the brain is essential for induction of pulmonary dysfunction and mortality in C-IRIS mice(Kawano et al., 2023b), we evaluated numbers of CD4^+^ T, Th1, and Th17 cell in the brains from mice treated with sGA. As expected, sGA treatment significantly reduced CD4^+^ T, Th1, and Th17cell numbers in the brain at 5 days after CD4^+^ T cell transfer in CnH99-infected mice (**Fig. 3B**). As we demonstrated in a previous report, microglia are upregulated in the C-IRIS mice(Kawano et al., 2023b). Here, we found that the microglia population was significantly suppressed by sGA treatment (**Fig. 3B**), suggesting sGA prevents neuroinflammation in the brains of C-IRIS mice. Together, these findings indicate that sGA suppresses C-IRIS disease by reducing CD4^+^ T/ Th1 cell proliferation in the lung, their infiltration into the brain, and their microglia activation in the brain.

**Figure 3.**
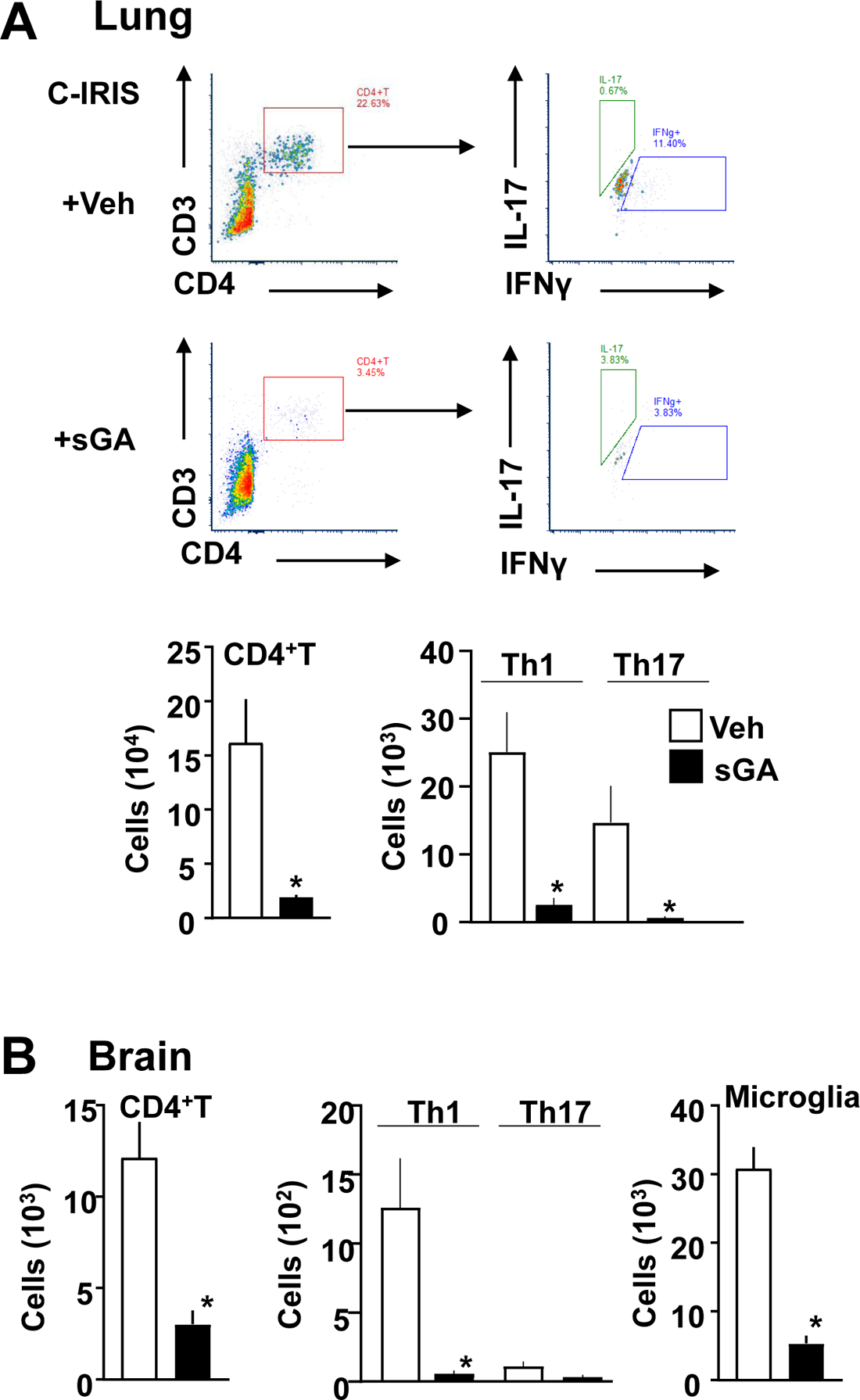
sGA reduces inflammatory cell populations in the lungs and brains of C-IRIS mice. Flow cytometry analysis demonstrated that sGA treatment significantly reduced the numbers of CD4+ T cells, Th1, and Th17 cells in the lungs (A) and brains (B) of C-IRIS mice. sGA also reduced microglia populations in the brain, indicative of suppressed neuroinflammation. Data are shown as mean ± SD (*p < 0.05).

### sGA ameliorated neuronal damage in the brain

Murine C-IRIS condition is associated with neurodegeneration due to the infiltration of CD4^+^ T cells into the brain tissue, which has neurotoxic properties(Kawano et al., 2023b), resulting in neurogenic pulmonary dysfunction. In addition, C-IRIS patients have multiple neurological symptoms, including headache, fever, cranial neuropathy, alteration of consciousness, lethargy, memory loss, meningeal irritation signs, and visual disturbance (Wu et al., 2020), likely mediated by neurodegeneration in the brain. Thus, we evaluated neuron numbers in various brain regions with potential involvement in neurological symptoms of C-IRIS. In addition, we evaluated neuronal status in the brain of C-IRIS mice after sGA treatment. As shown in (**Fig. 4**), Golgi cox-stained neuron numbers were significantly reduced in the prefrontal cortex (PFC), The basolateral amygdala (BLA), lateral hypothalamus (LH), and periaqueductal gray (PAG) of the C-IRIS mice. Following sGA treatment at 1 and 3 days after CD4^+^T cell transfer into CnH99-infected mice, the decrease in neuron numbers showed significant improvement in PFC, LH, and PAG (**Fig. 4**). Improved neuronal survival further supports the efficacy of sGA as a potential treatment for the prevention of C-IRIS induction.

**Figure 4.**
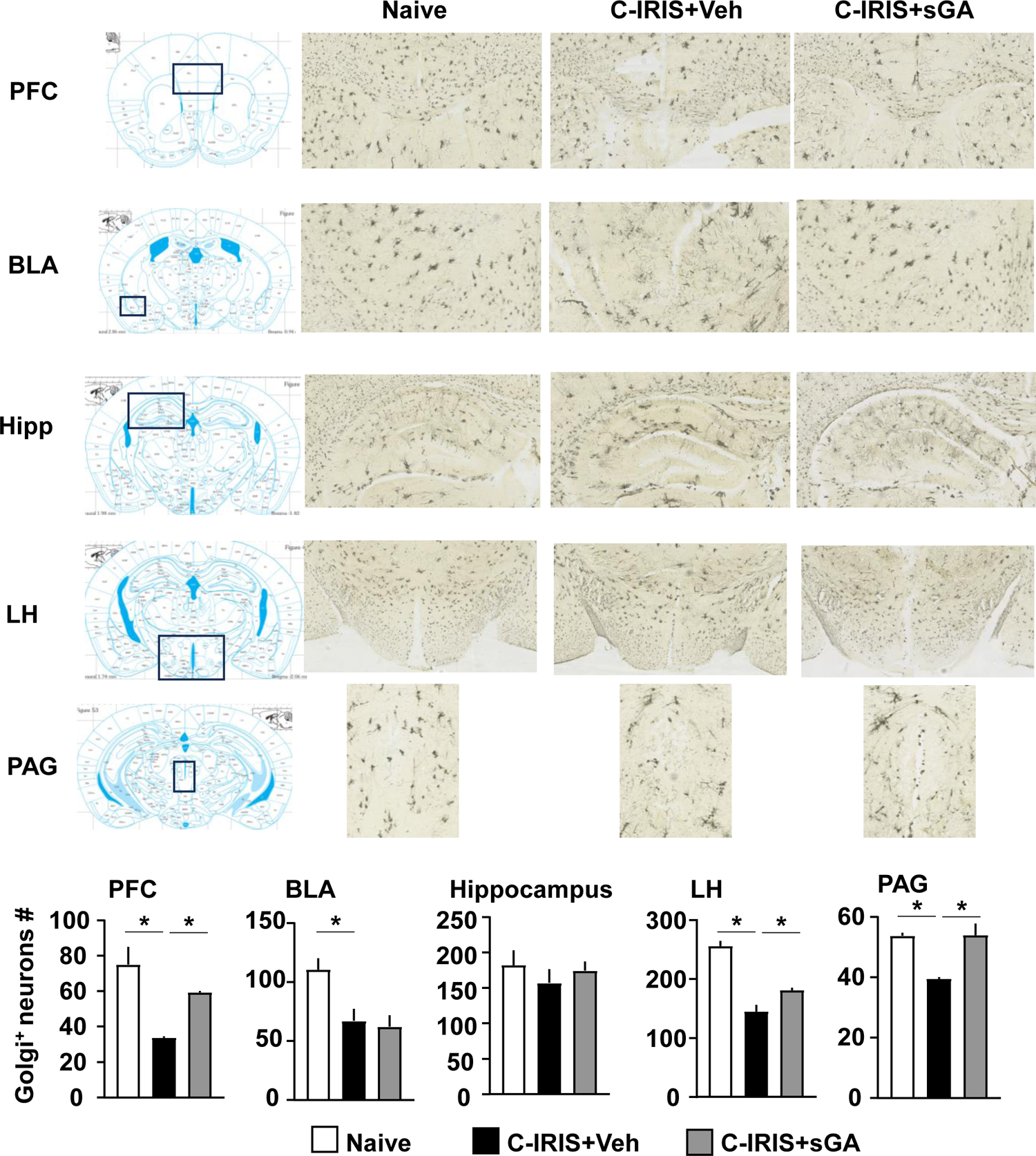
sGA mitigates neuronal damage in C-IRIS mice. Golgi-Cox staining revealed a significant reduction in neuron numbers in the prefrontal cortex (PFC),lateral hypothalamus (LH), and periaqueductal gray (PAG) in C-IRIS mice compared to naive controls. sGA treatment significantly protected against neuronal loss in these regions. Data are expressed as mean ± SD (*p < 0.05).

## Discussion

NSAIDs and corticosteroids are prescribed to suppress excessive inflammation in C-IRIS patients. However, such immunosuppressive medications may impair the immune response to existing infections and render patients more vulnerable to new infections(Meintjes et al., 2012b). Preventing CD4+ T cell migration to the brain and inhibiting Th1 differentiation could serve as promising therapeutic approaches for C-IRIS management.. In the present study, we investigated the effects of sGA, a polypeptide-based FDA-approved drug for treating MS, on an experimentally induced murine model of C-IRIS. Our principal findings are that sGA improved pulmonary dysfunction, reduced mortality, and prevented neuronal loss in the brain. Various studies have demonstrated that GA exerts its immunomodulatory effects by altering T-cell differentiation and promoting the proliferation of Th2-polarized GA-reactive CD4^+^ T cells(Makar et al., 2023). Consistent with the previous reports, we found that sGA treatment significantly reduces CD4^+^ T, Th1, and Th17 cell populations in the lung and the brain. Further, we observed increased numbers of microglia in the brain of C-IRIS mice and showed that the administration of sGA significantly downregulated the microglia activation. Evidence indicates that GA inhibits particular microglial receptors, such as the purinergic P2X7 ionotropic receptor (P2X7R), which is increased during inflammatory conditions(Aharoni, 2013). P2X7R is a receptor expressed on monocytes and microglia, and it plays a vital role in the activation and proliferation of microglia, which may cause tissue damage and neuroinflammation(Kasindi et al., 2022)—suggesting that the inhibitory effect of sGA on microglia could contribute to neuroprotection. It is important to note that unlike most Disease Modifying Therapies (DMTs) (Kasindi et al., 2022), sGA does not appear to suppress the peripheral immune response as most DMTs do. Therefore, sGA is likely to have minimal to no immunosuppressive effect in C-IRIS patients, thereby lowering the risk of secondary infection.

C-IRIS can present with a variety of clinical signs, including altered mental status personality and behavioral changes, confusion, hallucinations, and, in rare cases, lethargy(Brienze et al., 2021). The PFC is a part of the cerebral cortex responsible for many aspects of human behavior and is considered the “personality center” of the brain(Courtney, 2010). We observed a significant reduction of the neurons within the PFC of the C-IRIS mice, suggesting that neurodegeneration in the PFC could be responsible for the behavioral changes observed in C-IRIS patients is a critical region in the propagation and modulation of pain(Benarroch, 2012). Neuronal damage in the PAG of C-IRIS mice may be involved in chronic pain in C-IRIS patients. Further investigation warrants whether sGA treatment, due to its ability to reduce neuronal loss in PFC and PAG as seen in C-IRIS mice, could potentially preserve mental status and memory and prevent chronic pain in C-IRIS patients.

## Conclusion

In conclusion, sGA suppresses C-IRIS-induced Th1 and Th17 differentiation in the lung tissues and reduces CD4^+^ T, Th1, Th17, and microglia populations in the brain, resulting in improved respiratory dysfunctions, lowered mortality rate, and reduced brain neurodegeneration. These results suggest that sGA may be an effective therapeutic for alleviating C-IRIS disease. C-IRIS patients exhibit diverse clinical presentations, including headache, fever, cranial neuropathy, alteration of consciousness, lethargy, altered mental status, memory loss, meningeal irritation signs, and visual disturbance. Therefore, we will investigate the association of the aforementioned clinical signs with neurodegeneration within different brain regions and test the therapeutic efficacy of the sGA drug in improving these symptoms.

## Ethics statement

All mouse experiments were approved by the University of Illinois Animal Care and Usage Committee (IACUC) under protocol number 22140.

## Data Availability

The data supporting the findings of this study are available from the corresponding author upon reasonable request. Due to privacy and confidentiality considerations, the data are stored on our lab server and are not publicly available, but they will be shared with qualified researchers upon request.

## CRediT authorship contribution statement

S.A., J.Z., and M.I. planned the experiments and wrote the manuscript. S.A. and J.S. performed sample isolation. Z.S. and J.C. performed fungal burden studies. M.I. and K.S. planned and performed pulmonary function tests. L.K. performed Golgi-stained neuron counting.

## Acknowledgments

We thank Mary Clutter and Yee Ming Khaw for helping with sample isolation and analysis. This research was supported by NIH R01-AI136999 (MI). The authors declare no competing financial interests.

